# DeepCell Kiosk: Scaling deep learning-enabled cellular image analysis with Kubernetes

**DOI:** 10.1101/505032

**Authors:** Dylan Bannon, Erick Moen, Morgan Schwartz, Enrico Borba, Takamasa Kudo, Noah Greenwald, Vibha Vijayakumar, Brian Chang, Edward Pao, Erik Osterman, William Graf, David Van Valen

## Abstract

Deep learning is transforming the analysis of biological images but applying these models to large datasets remains challenging. Here we describe the DeepCell Kiosk, cloud-native software that dynamically scales deep learning workflows to accommodate large imaging datasets. To demonstrate the scalability and affordability of this software, we identified cell nuclei in 10^6^ 1-megapixel images in ~5.5 h for ~$250, with a sub-$100 cost achievable depending on cluster configuration. The DeepCell Kiosk can be downloaded at https://github.com/vanvalenlab/kiosk-console; a persistent deployment is available at https://deepcell.org.

## Main Text

While deep learning is an increasingly popular approach to extracting quantitative information from biological images, its limitations significantly hinder its widespread adoption. Chief among these limitations are the requirements for expansive sets of training data and computational resources. Here, we sought to overcome the latter limitation. While deep learning methods have remarkable accuracy for a range of image-analysis tasks including classification^1^, segmentation^2–4^, and object tracking^5,6^, they have limited throughput even with GPU acceleration. For example, even when running segmentation models on a GPU, typical inference speeds on megapixel-scale images are in the range of 5-10 frames per second, limiting the scope of analyses that can be performed on images in a timely fashion. The necessary domain knowledge and associated costs of GPUs pose further barriers to entry, although recent software packages^7–11^ have attempted to solve these two issues. While cloud computing has proven effective for other data types^12–15^, scaling analyses to large imaging datasets in the cloud while constraining costs is a considerable challenge.

To meet this need, here we have developed the DeepCell Kiosk (Fig. 1a). This software package takes in configuration details (user authentication, GPU type, etc.) and creates a cluster in the cloud that runs predefined deep learning-enabled image-analysis pipelines. This cluster is managed by Kubernetes, an open-source framework for running software containers (software that is bundled with its dependencies so it can be run as an isolated process) across a group of servers. An alternative way to view Kubernetes is as an operating system for cloud computing. Data is submitted to the cluster through either a web-based front-end, a command line tool, or an ImageJ plugin. Once submitted, it is placed in a database where the specified image-analysis pipeline can pick up the dataset, perform the desired analysis, and make the results available for download. Results can be visualized by a variety of visualization software tools^16,17^.

**Figure 1:**
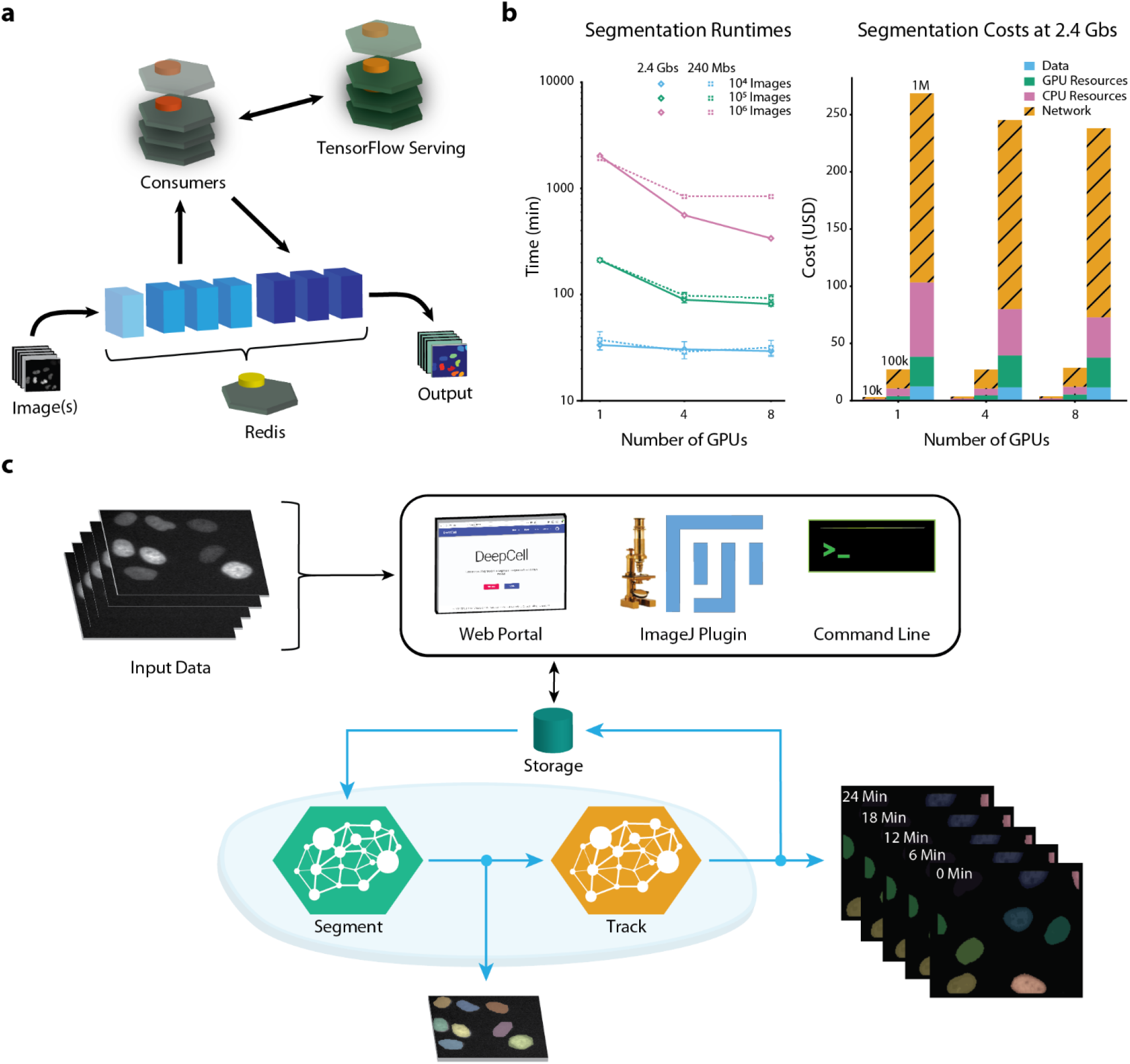
Architecture and performance of the DeepCell Kiosk. **a)** Data flow through the DeepCell Kiosk. Images submitted to a running Kiosk cluster are entered into a queue. Once there, images are processed by objects called consumers, which execute all the computational operations in a given pipeline. Consumers perform most computations themselves but access deep learning by submitting data to TensorFlow Serving and capturing the result. This separation allows conventional operations and deep learning operations to occur on different types of nodes, which is essential for efficient resource allocation. After processing, the result is returned to the queue, ready to be downloaded by the user. **b)** Left, Benchmarking of inference speed demonstrates scaling to large imaging datasets (see SI for details). Throughput is ultimately limited by data transfer speed. Right, an analysis of cost demonstrates affordability. Network costs are incurred by inter-zone network traffic in multi-zone clusters, which are more stable and scale faster than their single-zone counterparts. This cost can be avoided when users configure a single-zone cluster. **c)** The DeepCell Kiosk enables construction and scaling of multi-model pipelines. For example, a live-cell imaging consumer can access deep learning models for both segmentation and tracking. The live-cell imaging consumer places the frames for a movie into the queue for a segmentation consumer, where they are processed in parallel. Once segmented, the images are processed by a tracking consumer to link cells together over time and to construct lineages. The results, which consist of label images of segmented and tracked cells as well as a JSON file describing mother-daughter relationships, are uploaded to a cloud bucket, from which they can be downloaded by the user.

To ensure that image-analysis pipelines can be run efficiently on this cluster, we made two software design choices. First, image-analysis pipelines access trained deep learning models through a centralized model server in the cluster. This strategy enables the cluster to efficiently allocate resources, as the various computational steps of a pipeline (deep learning and conventional computer-vision operations) are run on the most appropriate hardware. Second, the DeepCell Kiosk treats hardware as a resource that can be allocated dynamically, as opposed to a fixed resource. This conceptual shift means that cluster size scales to meet data analysis demand: small datasets lead to small clusters, while large datasets lead to large clusters. This feature, which has been previously demonstrated for other biological data types^13^, is made possible by our use of Kubernetes, which enables us to scale analyses to large datasets, reducing analysis time while containing costs (Fig. 1b). A full description of the DeepCell Kiosk’s software architecture, scaling policies, and benchmarking is provided in the Supplemental Information.

Given the wide range of image-analysis problems that deep learning can now solve, we designed the DeepCell Kiosk with the flexibility to work with arbitrary collections of deep learning models and image-analysis pipelines. Pipelines can exploit multiple deep learning models—and even other pipelines—while still leveraging the scaling ability provided by Kubernetes. For example, we constructed a pipeline (Fig. 1c) that pairs deep learning models for cell segmentation and cell tracking^5^ in order to analyze live-cell imaging data. Additional examples are described in the Supplemental Information, and documentation on creating custom pipelines is available at https://deepcell-kiosk.readthedocs.io.

In conclusion, the DeepCell Kiosk enables cost-effective scaling of customizable deep learning-enabled image-analysis pipelines to large imaging datasets. The dynamic nature of our scaling enables individual users to process large datasets. Further, it allows a single cluster to serve an entire community of researchers—much like BLAST^18^ has rendered sequence alignment accessible to the scientific community through a web portal. At the cost of network latency, software ecosystems like ImageJ can access complete deep learning-enabled pipelines hosted by such a cluster through a plug-in. While the data analyzed here came from *in vitro* cell culture, our emphasis on deep learning makes it possible for our software to analyze tissue data, including spatial genomics data^19^. We believe that by reducing costs and time-to-analysis, this work will accelerate the rate of biological discovery and change the relationship that biologists have with imaging data. Further, by focusing on deployment, this work highlights aspects of the performance of deep learning models beyond accuracy (such as inference speed) that significantly impact their utility. Finally, this work highlights the growing definition of the term ‘software.’ Just as deep learning has made data a part of the software stack, Kubernetes can do the same for hardware. Jointly developing data, code, and compute will significantly improve the performance of scientific software crafted in the era of Software 2.0.

## Acknowledgments

We thank numerous colleagues including Anima Anandkumar, Michael Angelo, Justin Bois, Ian Brown, Andrea Butkovic, Long Cai, Isabella Camplisson, Markus Covert, Michael Elowitz, Jeremy Freeman, Christopher Frick, Lea Geontoro, Andrew Ho, Kevin Huang, KC Huang, Greg Johnson, Leeat Keren, Daniel Litovitz, Derek Macklin, Uri Manor, Shivam Patel, Arjun Raj, Nicolas Pelaez Restrepo, Cole Pavelchek, Sheel Shah, and Matt Thomson for helpful discussions and contributing data. We gratefully acknowledge support from the Shurl and Kay Curci Foundation, the Rita Allen Foundation, the Paul Allen Family Foundation through the Allen Discovery Center at Stanford University, The Rosen Center for Bioengineering at Caltech, Google Research Cloud, Figure 8’s AI For Everyone award, and a subaward from NIH U24CA224309-01.

## Author Contributions

DB, WG, and DVV conceived the project; DB, WG, EO, and DVV designed the software architecture; DB, EO, and WG wrote the core components of the software; DB, EM, MS, EB, VV, BC, EO, WG, and DVV contributed to the code base; TK and EP collected data for annotation; EM, MS, NG, DB, WG, and DVV wrote documentation; DB, EM, WG, and DVV wrote the paper; DVV supervised the project.

## Supplemental Information

This Supplemental Information describes in further detail the software architecture of the DeepCell Kiosk, presents all our benchmarking data, and outlines the reasoning behind several of our design choices. Because we use terminology that is common in the cloud computing literature but may be unfamiliar to readers, we have included a glossary of common terms.

**Glossary**
Deep learning: A set of machine-learning techniques that learn effective representations of data with multiple levels of abstraction^1^.
TensorFlow: An open-source software library that is commonly used to build neural networks^2^.
Model Server: A web server that hosts deep learning models and returns predictions on data sent to the server.
TensorFlow Serving: Part of the TensorFlow tool ecosystem. A model server serving TensorFlow models.
Git: A version-control system that tracks changes in source code during software development^3^.
GitHub: A web-based platform that hosts code repositories under version control using Git^4^.
Container: A self-contained software unit consisting of all the code and dependencies necessary for quickly running an application; roughly equivalent to a virtual machine.
Docker: A program that creates containers and allocates computational resources to them^5,6^.
Dockerfile: A text document utilizing a Docker-specific markup language to specify the contents of a container.
Docker Image: A template for a container, built according to specifications in a Dockerfile.
Docker Container: A container run by Docker specifically. An instance of a Docker Image.
Microservice Architecture: A software architecture paradigm in which multiple independent processes communicate with one another to accomplish a larger task^7^.
REST API: An API model that transfers data using human-readable data structures and engages in recurring handshaking protocols.
gRPC API: An API model that entails transferring data using binary (non-human-readable) data structures and minimizing the number handshakes that occur.
Kubernetes: A system for coordinating communication between and replication of containers^8^. Kubernetes can facilitate a microservice architecture in which each container is a separate process.
Redis: An open-source in-memory database to store key-value pairs^9^.
Helm: A package manager for Kubernetes^10^.
Prometheus: An open-source monitoring framework for Kubernetes^11,12^.
Consumer: Programmatic object that executes all the computational steps of an image-analysis workflow.
Queue: A sequence of data items submitted for processing.
Horizontal Autoscaling: Increasing computational resources by using more nodes.
Vertical Autoscaling: Increasing computational resources by using more-powerful nodes.
Kubernetes – Pod: A group of one or more containers with shared storage/network and a unique network identity. A pod represents a single instance of an application in Kubernetes^13^.
Kubernetes – Replica: Identical copies of a Pod^13^.
Kubernetes – Node: A worker machine that can run Pods^13^.
Kubernetes - Node pool: A collection of Nodes within a cluster that have the same configuration^13^.
Kubernetes - Metrics server: A cluster wide aggregator of resource usage data^13^.
Google Compute Engine: Google’s pool of cloud resources.
Google Kubernetes Engine: Google’s managed Kubernetes service.

### DeepCell Kiosk Architecture Overview

The DeepCell Kiosk uses Kubernetes to create scalable compute clusters, comprised of Docker containers, on Google Compute Engine. A schematic of a representative DeepCell Kiosk cluster is shown in Figure S1.

**Figure S1:**
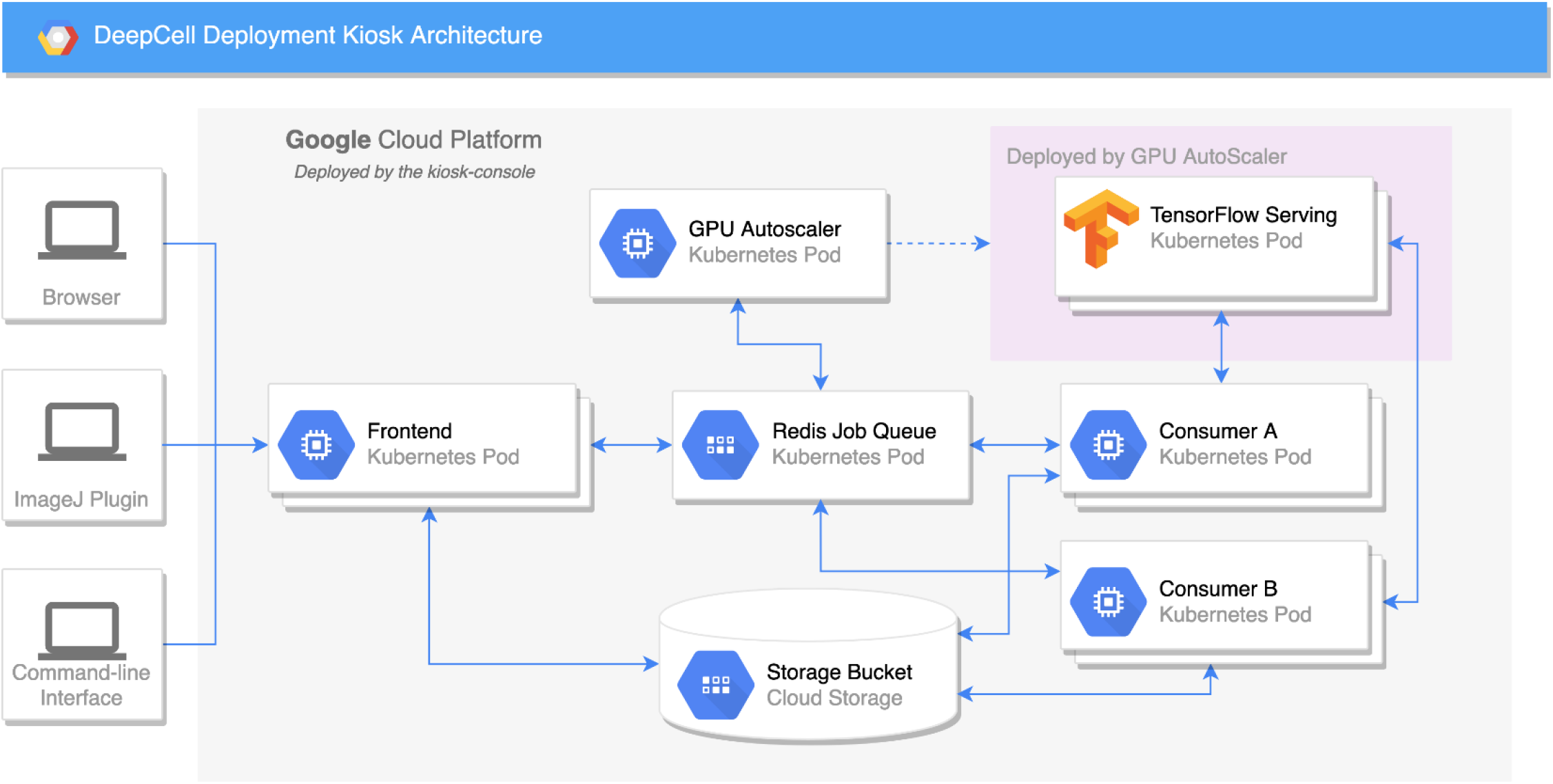
The software architecture of the DeepCell Kiosk and data flow within an operational cluster. Rectangles represent a replica set of 1 or more identical Kubernetes pods; cylinders represent cloud-provided storage space for data. All the pods in a given replica set perform the same logical function. Data enter through the “Frontend”, which uploads data into the “Storage Bucket” and creates an entry in the “Redis Job Queue”, which houses a queue of pending analysis jobs in a Redis database. The “GPU Autoscaler” monitors the “Redis Job Queue” and requisitions the first GPU node (pink rectangle) if there are pending jobs and no GPU nodes. Requisitioning of a GPU leads to creation of the “Tensorflow Serving” pod, which serves deep learning models for the entire cluster. The “Consumer” pods execute analysis workflows on images in their associated queue in “Redis Job Queue”. These pods access deep learning models by sending data back and forth to the “Tensorflow Serving” pod. Multiple types of consumers (e.g., A and B) can run simultaneously. Moreover, while each type of consumer has their own associated queue, they can access any queue present in Redis Job Queue. This allows consumers to interact with each other and facilitates construction of highly parallelized workflows. The output of an analysis workflow is placed in the “Storage Bucket”, from which it can be downloaded by the user.

### Software Design Decisions

DeepCell Kiosk users start the cluster-creation process by providing authorization credentials and details for the cluster configuration. These details include the GPU type (V100, T4, or P100) and the maximum allowed number of GPU nodes. Within the cluster, a microservice architecture is used to implement image-analysis workflows. Data enter the system through the front end and are placed in a queue (Fig. S1). Consumer objects (described in the “Consumers and Queues” section) manage the processing of each data item and access deep learning through calls to the model server. By effectively having two “services”—one that uses deep learning and one that does not— computational resources are efficiently allocated to meet demand. Further, Kubernetes enables this scaling to be dynamic, as the cluster can grow or shrink to meet demand.

Below, we describe a series of software design decisions that enabled the key features of the DeepCell Kiosk. Each of these design decisions was meant to facilitate three key features: *scalability, flexibility*, and *affordability*.

- <u>*Containerization*</u>. To facilitate the scalability, flexibility, and affordability of a DeepCell Kiosk cluster, we chose to containerize all code used in the project. See below for details on achieving scalability, which involves Kubernetes. For flexibility, having all code containerized means that users who wish to implement custom workflows need only modify well-defined components of the DeepCell Kiosk and can be confident that all other components will continue to work as intended. Cost-savings come about through our ability to only requisition expensive computing resources (GPUs, large amounts of RAM) for containers where it is necessary (e.g., TensorFlow Serving) and to run other containers using much more affordable computing resources. We follow the design principle of “one process per container” to enhance security and resilience while accelerating the development lifecycle.
- <u>*Container orchestration through Kubernetes*</u>. We use Kubernetes and its associated package manager, Helm, to manage the allocation of compute resources and the ad-hoc replication of containers in the DeepCell Kiosk. We also use Google Kubernetes Engine (GKE) to requisition compute resources. Kubernetes performs several key functions, including:

○ Organizing containers into functional units called pods (in our architecture most pods consist of one container);
○ Requisitioning compute nodes (both with and without GPUs) from the cloud (GKE);
○ Assigning the appropriate pods to each node;
○ Scaling the number of nodes to meet demand (GKE); and
○ Facilitating internal communication between containers within the deployment.
- <u>*Infrastructure as code*</u>. Cloud computing requires a precise specification of the computational resources needed from the cloud. In the case of the DeepCell Kiosk, these configuration details include the location of storage buckets and the numbers and types of compute nodes in an active cluster. The configuration of each type of compute node requires more information, such as CPU type, memory size, the presence/absence of a GPU or other hardware accelerator, and the maximum allowable number of these nodes in a cluster. We present these choices to the user via a configuration menu with reasonable default settings. The specification for a cluster gets saved in environmental variables, which are sent to Google Cloud for cluster creation. They are also used to configure YAML files that specify which nodes each pod should occupy. In this fashion, we treat the hardware running our cluster as software, a concept called “infrastructure as code”. This strategy enables us to treat deep learning models and deployment systems collectively as software.
- *<u>Preemptible instances</u>*. To decrease the cost of analyzing large datasets, we configured the DeepCell Kiosk to request preemptible GPU nodes. While preemptible nodes can be interrupted, they are ~3 times cheaper than regular instances^14^ and are suitable for large inference tasks. Further, we have engineered the DeepCell Kiosk to be tolerant of unanticipated node shutdowns, so that even if work is lost because a node suddenly shuts down, that work is guaranteed to be repeated on another node.
- *<u>Logging and monitoring</u>*. We use Prometheus to monitor cluster performance metrics and the ELK stack (Elasticsearch, Logstash, and Kibana) to enable logging throughout the Kubernetes cluster. Robust logging facilitates development of custom data-processing pipelines, while metric collection provides the real-time data needed for horizontal pod autoscaling. Additionally, these metrics enable benchmarking of the performance and cost of new pipelines.
- *<u>Inter-service communication with gRPC</u>*. Because large amounts of imaging data are shuttled between two services during analysis tasks, the communication between services needs to be efficient. REST APIs are common communication paradigm that represent data in a human-readable format. Unfortunately, they were not designed for efficiency. Because the DeepCell Kiosk was designed to maximize data throughput, we use gRPC as an alternative, more efficient means of communicating between services in the cluster.
- *<u>Continuous integration/continuous delivery</u>*. We use Travis CI to implement continuous integration/continuous delivery. This online service allows changes to the code base to be reflected in a deployment, provided the modifications pass the existing unit tests. Developers can therefore rapidly implement new features and deploy novel workflows in the DeepCell Kiosk.
- *<u>User interface</u>*. We have created a simple drag-and-drop user interface in JavaScript using the libraries React, Babel, and Webpack. This interface allows users to submit data to the Kubernetes cluster via a web browser. Alternatively, users can submit data to the cluster directly using Python code, or via the DeepCell ImageJ plugin. Together, these features render the cluster accessible and help flatten the learning curve for inexperienced users.

### Consumers and Queues

Conceptually, the DeepCell Kiosk relies on two objects for scaling image analysis workflows: consumers and queues (Fig. S2). Consumers contain Python code that specifies a sequence of computations that must be performed on an image or collection of images for a given workflow. Consumers are similar to “workers” in parallel computing schema; we use the term “consumer” here due to our use of the Redis database. For example, a segmentation consumer that relies on a deep watershed algorithm might specify the following steps:

1. Contrast adjust and normalize the input image.
2. Process the image with a deep learning model to create a prediction of the distance map.
3. Threshold the distance map to create markers for each object.
4. Perform marker-based watershed to produce the final label image for the input image.

To execute these steps, the consumer retrieves the input image from a queue, performs all the specified operations in sequence, and returns the final label image to the user. Note that all operations in the pipeline take place within the consumer’s node, except operations involving deep learning. These operations are performed by submitting requests to TensorFlow Serving pods, which exist on nodes with GPUs. After receiving the model output, the consumer commences with the rest of its processing steps.

**Figure S2:**
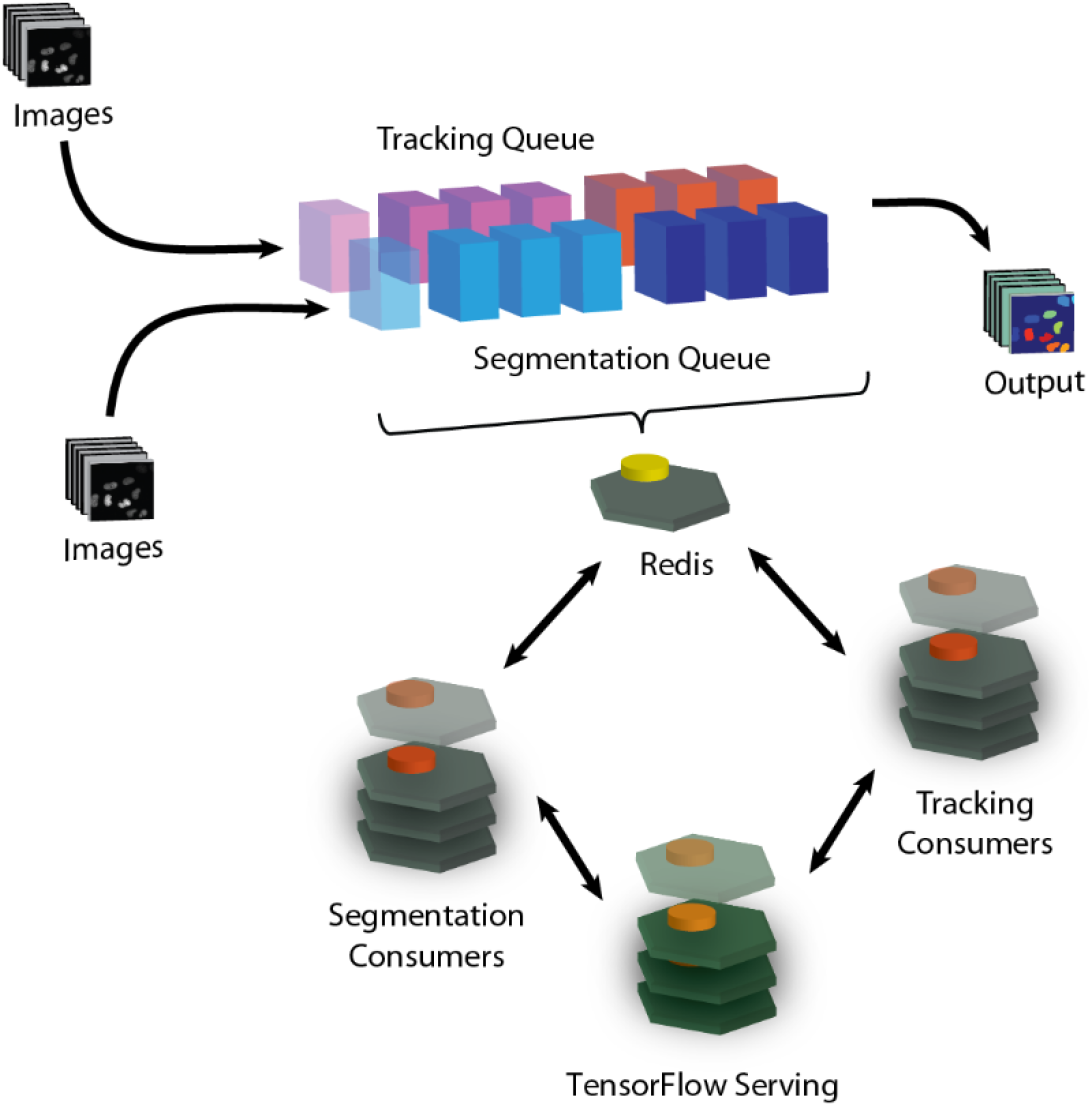
Consumers and queues. Consumers process data items with defined computational workflows, while queues contain links to data items to be processed by consumers. Each consumer type has an associated queue. The number of active consumers of a given type dynamically scales with analysis demand.

Why not just add a GPU to the consumer? The answer is cost efficiency. By separating deep learning and non-deep learning operations into separate pods, we can efficiently allocate resources as the analysis scales. For instance, take the case where the non-deep learning operations in a pipeline take twice as long to execute as the deep learning operations. If all the operations occurred sequentially on the same node, then two-thirds of the GPU cost would be wasted because the GPUs remain idle during non-deep learning operations; throughput for each GPU would be limited as well. Separating these two operations into their own pods significantly increases the cost efficiency and hence scalability.

We note that this approach relies on a low cost of data transfer between consumers and TensorFlow Serving. For single-zone clusters (in which all nodes exist within the same geographic location), there is no cost to transfer data between nodes. For multi-zone clusters, which are more stable, nodes can exist in different zones and network ingress fees for data transfer between nodes in different zones apply. An analysis of this extra cost is given in Figure 1 of the main text; for image segmentation, we find that this extra cost can more than double the operating cost of a cluster. For this reason, the default setting is to create single zone clusters.

The default configuration of the DeepCell Kiosk is linked to our lab’s collection of consumers and models. At the time of publication, these consumers—and the associated models—include:

- Segmentation consumer: segments nuclei in images of cells in cell culture and tissues with a deep watershed model, as well as brightfield and fluorescent cytoplasm images of cells in cell culture.
- Tracking consumer: segments and tracks single cells in live-cell imaging movies.

For users who wish to construct their own consumers to be used with their own deployment of the DeepCell Kiosk, instructions can be found at https://deepcell-kiosk.readthedocs.io/en/master/index.html.

### Horizontal Pod Autoscaling

Scaling computational resources to meet analysis demand while minimizing costs is one of the key features of the DeepCell Kiosk. Conceptually, there are two ways to increase computing resources in a cluster. In the first way, one can create more pods and provide each pod with access to the same amount of CPU power and memory as the existing pods (horizontal pod autoscaling). Alternatively, the number of pods can be held fixed and the amount of CPU and/or memory available to each pod can be increased (vertical autoscaling). We have, in all instances, opted for horizontal autoscaling over vertical autoscaling, ultimately because our use case (identically processing many, many images of identical size) is highly parallelizable at the image level.

Note that scaling in the DeepCell Kiosk is constantly occurring on two levels. First, scaling occurs at the pod level, where Kubernetes is determining whether the current numbers of pods are appropriate for the current amount of work in the cluster. Second, Google Kubernetes Engine is determining whether there are enough computational resources already allocated to the cluster; if not, then the number of nodes is scaled up or down.

The DeepCell Kiosk’s approach to scaling focuses on optimally utilizing the cluster’s most expensive resource: GPUs. If demand is high, then additional GPUs need to be recruited; if demand is low, then the number of GPUs needs to be reduced. Additionally, if there are no data to be analyzed for an extended period of time, then the number of GPUs should be reduced to zero. Accordingly, the DeepCell Kiosk has two phases for scaling: 0-to-1 and 1-to-many. The details of these phases are described below. For both phases, the cluster makes use of metrics provided by Prometheus.

When the cluster is first created or when it is inactive for a long period of time (~5 min), the number of GPUs attached to the cluster is 0. When a new data item enters a queue, a GPU needs to be requisitioned for it to be processed. This step is referred to as 0-to-1 scaling, and the ability to perform 0-to-1 scaling is what enables us to turn GPUs off when they are not being used. Because performing 0-to-1 scaling is not an ability that is native to Kubernetes, we developed a custom Kubernetes pod that performs this task. This pod is created upon cluster construction and remains active during a cluster’s lifecycle.

Once a GPU has been requisitioned, data can be processed with deep learning-enabled consumers. As more data enter the system, scaling is necessary to meet the demand for analysis, including scaling the number of active consumers to control how many data items are analyzed in parallel. Scaling also addresses underlying computational resources—both CPUs and GPUs. This phase is referred to as 1-to-many scaling, for which we use the horizontal pod autoscaling system internal to Kubernetes (Fig. S3).

The system that Kubernetes uses for horizontal pod autoscaling is detailed in the Kubernetes online documentation^15^. Briefly, for each pod, one defines a metric and a target value for that metric. The metric typically represents the utilization of a resource, for example a GPU. If the metric (utilization) is high, then more resources are needed, and the pod is scaled up. If the metric is low, then fewer resources are needed and the pod is scaled down. Scaling the pod up or down often requires the addition or subtraction of compute nodes, which is performed by Kubernetes. Every 15s the cluster evaluates the metrics to determine the desired number of pods:

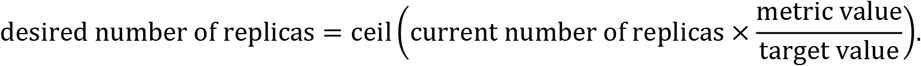

The value of the metric used in this formula is its average over this time period.

We define two classes of metrics, one for consumers (recall that each type of consumer is its own pod) and one for TensorFlow Serving. Recall that consumers exist on CPU-only nodes, while TensorFlow Serving exists on GPU nodes. Our desired behavior is:

- If GPU utilization is high because of high analysis demand, then increase the number of TensorFlow Serving pods and hence the number of GPUs.
- If there is a high analysis demand (there are a lot of data items in the consumer’s queue), then increase the number of consumers.
- If GPU utilization is too high (e.g., ~100%), then reduce the number of consumers to prevent an internal denial of service attack on the TensorFlow Serving pods.

**Figure S3:**
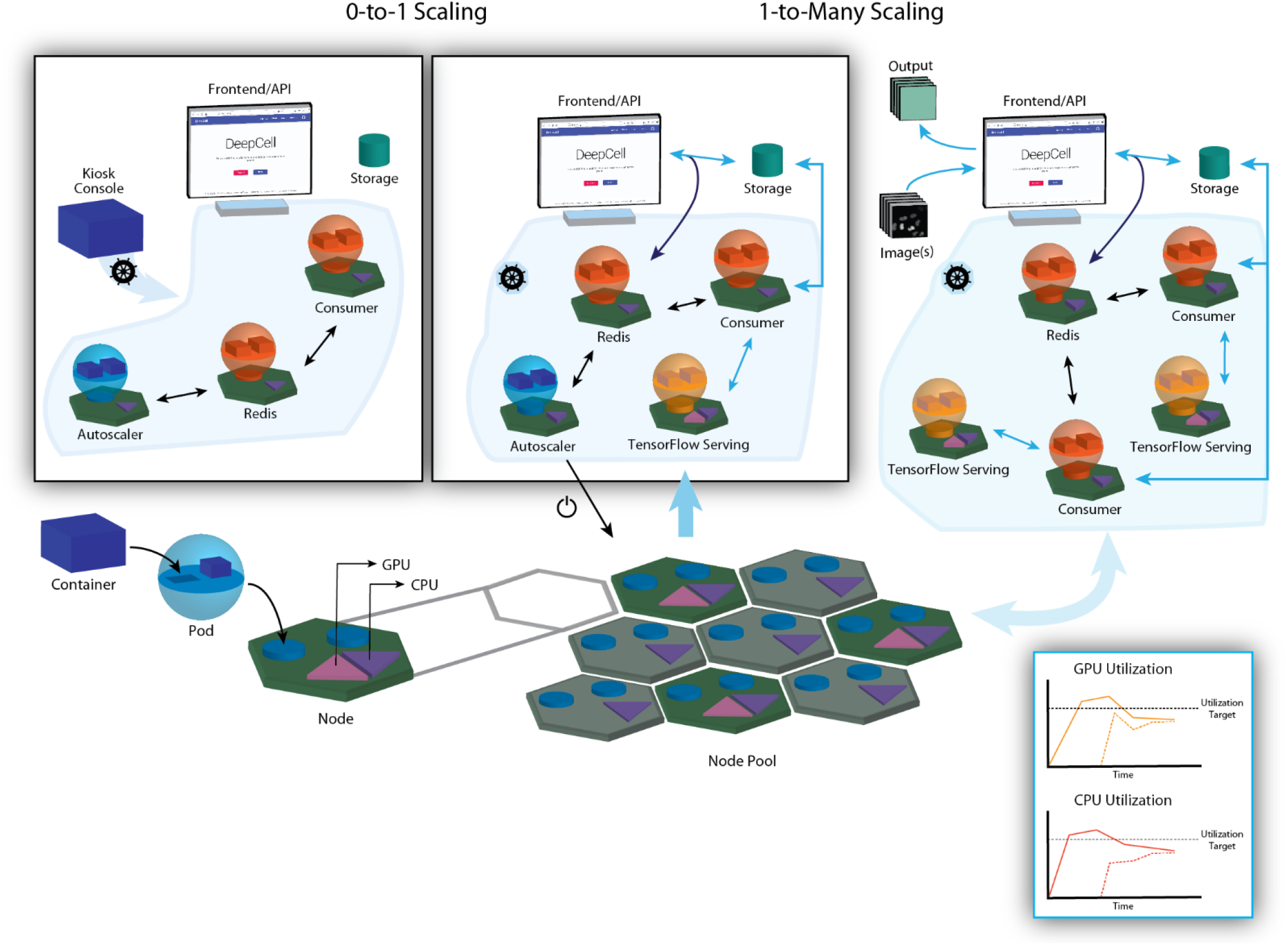
Horizontal pod autoscaling with Kubernetes. Each pod is given a metric and a target. The metric is averaged over a given time period (15 s in the default configuration of the DeepCell Kiosk), and a comparison of this average value with the target value determines whether a pod will be scaled up or down. If a metric is greater than its target, then the pod will be scaled up; if less, then it will be scaled down. If the metric and target are within a certain tolerance (The DeepCell Kiosk uses 10%), then no scaling occurs.

We have found that the following definitions of metrics and targets lead to robust and stable scaling:

- For TensorFlow Serving, we define a simple metric and constant target value for that metric:

○ Metric = Percent GPU utilization (0-100)
○ Target= 70
- For consumers, we define a more complex metric with three regimes and a constant target value:

○ No work: Metric = 0 if the number of TensorFlow Serving pods is 0 and there are 0 data items in the queue
○ Some work:

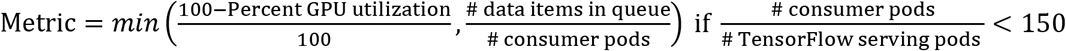
○ Too much work: Metric = 0.1125 otherwise
○ Target = 0.15.

These definitions for metrics and targets provide the desired behaviors, as demonstrated in the main text. The definitions for the TensorFlow Serving pod seek to maintain GPU utilization at 70%.

For the consumer pods, the three metric regimes coordinate to produce the desired behavior. In the “no-work” metric regime, there is no work to do, and we ensure that there is no scaling up by pinning the metric value to 0.

The “some work” metric regime uses a min() function to reliably scale the cluster up and down. During scale up, the first term in the min() function is smaller, which ties the number of consumers to GPU utilization. Recall that GPU utilization is a proxy for the activity of TensorFlow Serving. This term tells the system to increase the number of consumers until TensorFlow Serving is driven to the point that GPU utilization is 85%. Because the value of GPU utilization that triggers the scale up of TensorFlow Serving is 70%, this increase in consumers effectively forces the system to overshoot the target value and to keep requesting more resources indefinitely until either the Google Cloud account’s limit for GPUs is reached or the work is finished.

When it comes to consumer scale-down in the “some work” metric regime, a mechanism unrelated to GPU utilization is needed; note that any consumer type can drive GPU utilization, even in the absence of work for other consumer types. This mechanism is furnished by the second term in the min() function. As the consumer’s queue empties, this term drops below the value of the first min() term and then pushes the number of consumer pods down to zero as the queue becomes progressively emptier.

Note that the “some work” metric regime only holds when (# consumer pods / #TensorFlow serving pods) < 150. We add this condition because we have empirically found that for models with inference speeds of 5-10 images/s, having this ratio higher than 150 substantially increases the risk of an internal denial of service attack on the TensorFlow Serving pod (data not shown). When this condition no longer holds, perhaps due to transiently overzealous scaling up of consumer pods, we enter the “too much work” metric regime, where we pin the value of the metric at 0.1125, 75% of the target value of 0.15, ensuring a consistent and moderately-paced scale down of the consumer pods.

The target value of the consumer autoscaling metric and many of the metric’s parameters were determined heuristically using our single-cell segmentation models, and we have found that they support scaling for a modest variety of model types and multi-model workflows. However, given the heuristic origin of the parameters’ values, customization may be necessary as the number of available models and workflows expands.

### Cluster Scaling Analysis

Data transfer poses a fundamental limit to how much data can be analyzed in the cloud. Given the high data transfer speeds between nodes and storage databases within a cloud computing system, the biggest bottleneck to analysis is generally the transfer of data from a local workstation into a cloud bucket. A cluster that is operating efficiently will process data at the same rate it enters the cluster.

Given a data transfer speed *d* and a single model with an inference speed *s*, we can write the relation

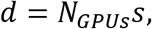

where *N_GPUs_* is the maximum useful number of GPUs in the cluster.

For cost-savings, the default maximum number of GPUs in a cluster is relatively small. This means that, often, the GPU-dependent deep learning step will be the rate-limiting step in an image processing pipeline. However, this also means that the way to optimize cluster performance is to have access to a sufficiently large number of GPUs, given data upload speed and image processing speed. To aid users in optimally setting the maximum number of GPUs the DeepCell Kiosk should have access to, we have included Figure 4, which plots this relationship for reasonable values of the parameters. Note that users can add GPUs beyond the maximum useful number, but these extra GPUs would increase costs and produce no tangible increase in image processing speed.

**Figure S4:**
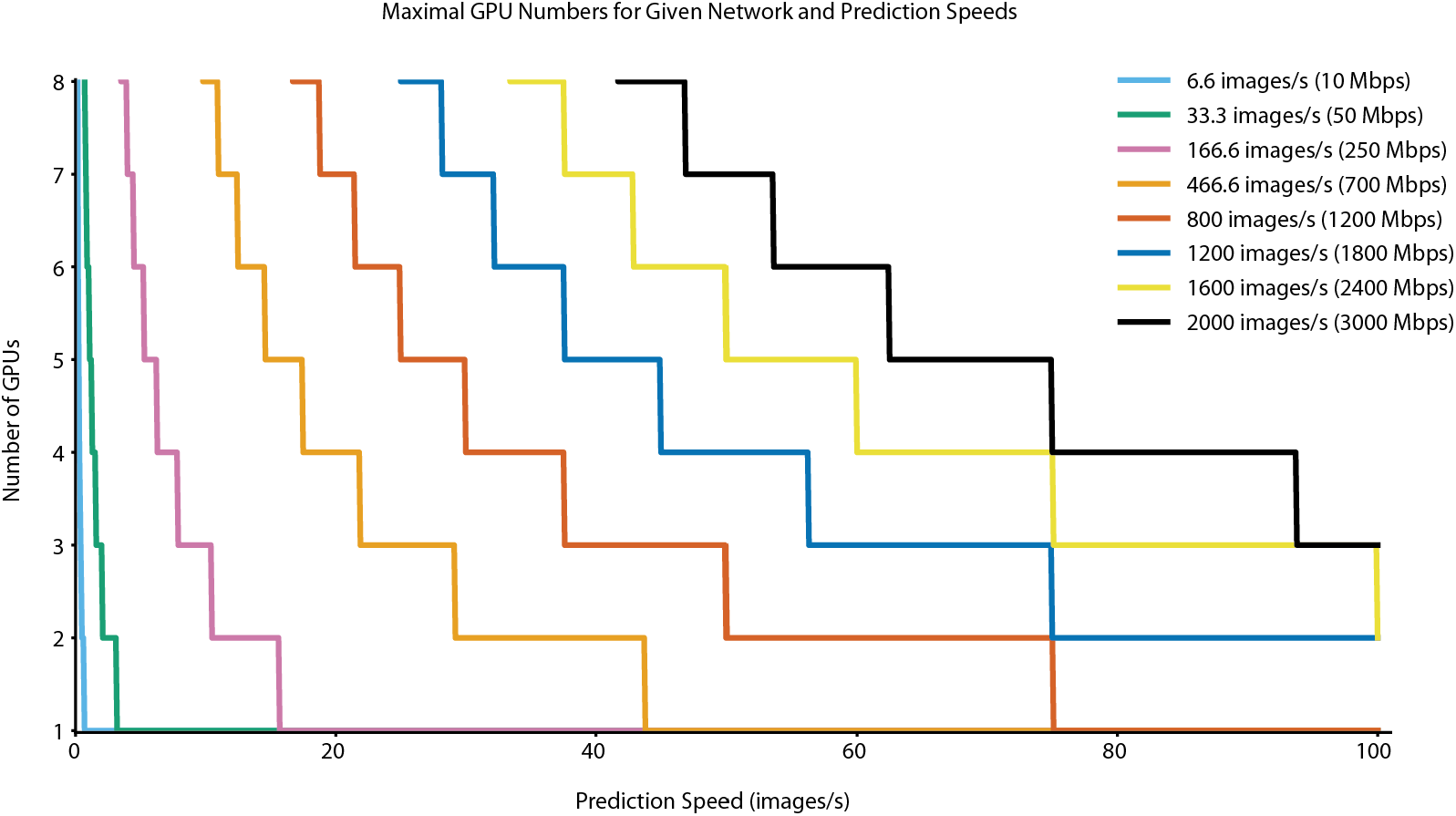
The relationship between data upload speed, model inference speed, and number of GPUs in a cluster. Each line represents a solution to the above equation where the rate of image processing in a cluster is balanced with the rate of image upload to a cluster. On one hand, faster models require fewer GPUs to process images at any given upload rate. On the other hand, faster data upload speeds necessitate the presence of more GPUs in order to process images as quickly as they’re being uploaded. Data transfer speed and model speed combined determine the maximal useful number of GPUs for a cluster. This figure assumes that all images are 1-megapixel images that occupy approximately 1.5 Mb on disk.

### Kiosk Benchmarking

We created a utility to benchmark the scalability and performance of the DeepCell Kiosk. Instructions for using this utility and creating the figures in this paper are described at https://github.com/vanvalenlab/publication-figures/tree/master/2020-Bannon_et_al-Kiosk.

The workflow we benchmarked was a deep watershed nuclear segmentation workflow; training of the model used in this workflow is described in the “Model Architecture and Training” section of this supplement. To benchmark scalability and performance, we chose 100 images from our training data at random and combined them into a zip file. This file was then submitted to an active cluster a predetermined number of times. This process was repeated in triplicate for each condition, except for 1,000,000 image runs, which were run a single time.

The benchmarking utility records processing times and cluster size over time during inference; these are used to compute the total processing time and cluster cost for a given benchmarking job. The total compute cost for a benchmarking job is the number of node-hours for each node type multiplied by the hourly cost for that node type, summed over all the different node types. This is given by

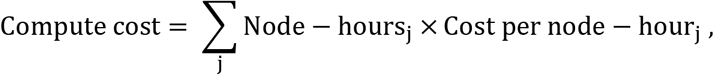

where j indexes the different node types. Costs for each type of node (current as of 06/17/2019) are defined in https://github.com/vanvalenlab/kiosk-client/blob/master/kiosk_client/cost.py

Because we wish to account for network fees, we tracked the transfer of data during workflow inference. Data transfer can be broken into the following 10 steps.

1. Upload of zip file to storage bucket
2. Download of zip file from storage bucket by zip-consumer pod
3. Upload of individual raw images to storage bucket by zip-consumer pod
4. Download of individual raw images from storage bucket by redis-consumer pod
5. Transfer of data to TensorFlow-Serving pod by redis-consumer pod
6. Transfer of model inference results to redis-consumer pod by TensorFlow-Serving pod
7. Upload of individual predicted image to storage bucket by redis-consumer pod
8. Download of individual predicted image from storage bucket by zip-consumer pod
9. Upload of zipped predicted images to storage bucket by zip-consumer pod
10. Download of zipped predicted images from storage bucket by client

Using this breakdown, we then computed 4 classes of data storage and network fees, which are described below.

- General storage fee: This is the cost of storing data in a cloud bucket
- Storage network egress fee. We assume that the user is in a different zone than the cloud bucket, and hence there is an egress fee for files downloaded by the client.
- Storage operations fee: File operations (e.g., construction, deletion, etc.) in cloud buckets incur a fee
- Inter-zone egress fee: If a cluster has a multi-zone configuration, there is no guarantee that redis-consumer pods and TensorFlow serving pods will be in the same zone. If they are in different zones, there is a network egress fee for data sent back and forth between these two pods.

Fees were calculated using the formula provided by Google Cloud (https://cloud.google.com/compute/all-pricing#network_pricing); the full implementation can be viewed at https://github.com/vanvalenlab/publication-figures/blob/master/2020-Bannon_et_al-Kiosk/figure_generation/data_extractor.py. The general storage, storage egress, and storage operations fees were reported as “Data” fees while the inter-zone egress fee was reported as “Network” fees in Figure 1. We again note that the inter-zone egress fees can be avoided by configuring the DeepCell Kiosk to create single-zone clusters.

**Figure S5:**
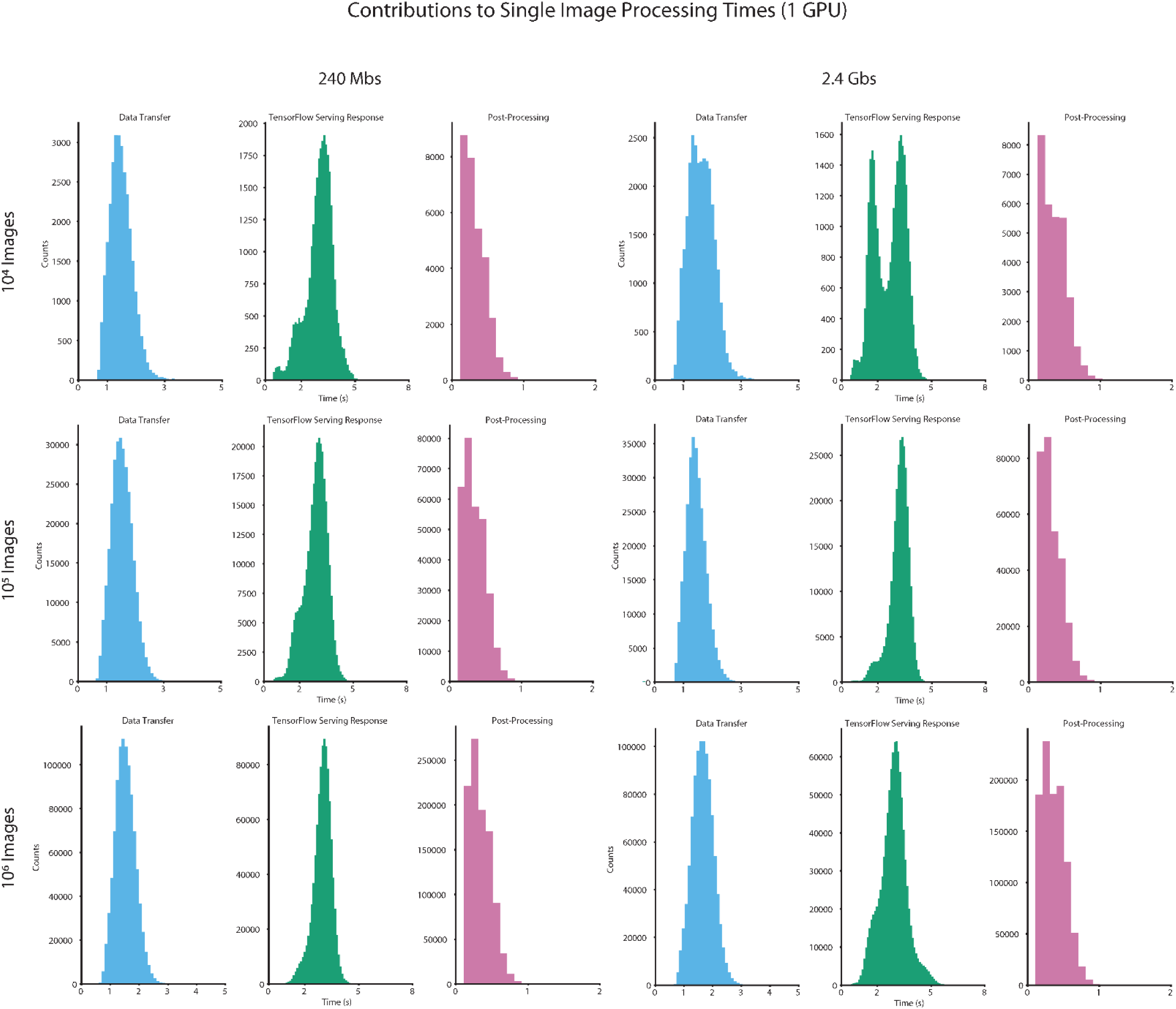
Benchmark timing data for 1 GPU clusters. Distributions of data transfer time, Tensorflow Serving response time, and postprocessing time required to process a single image.

**Figure S6:**
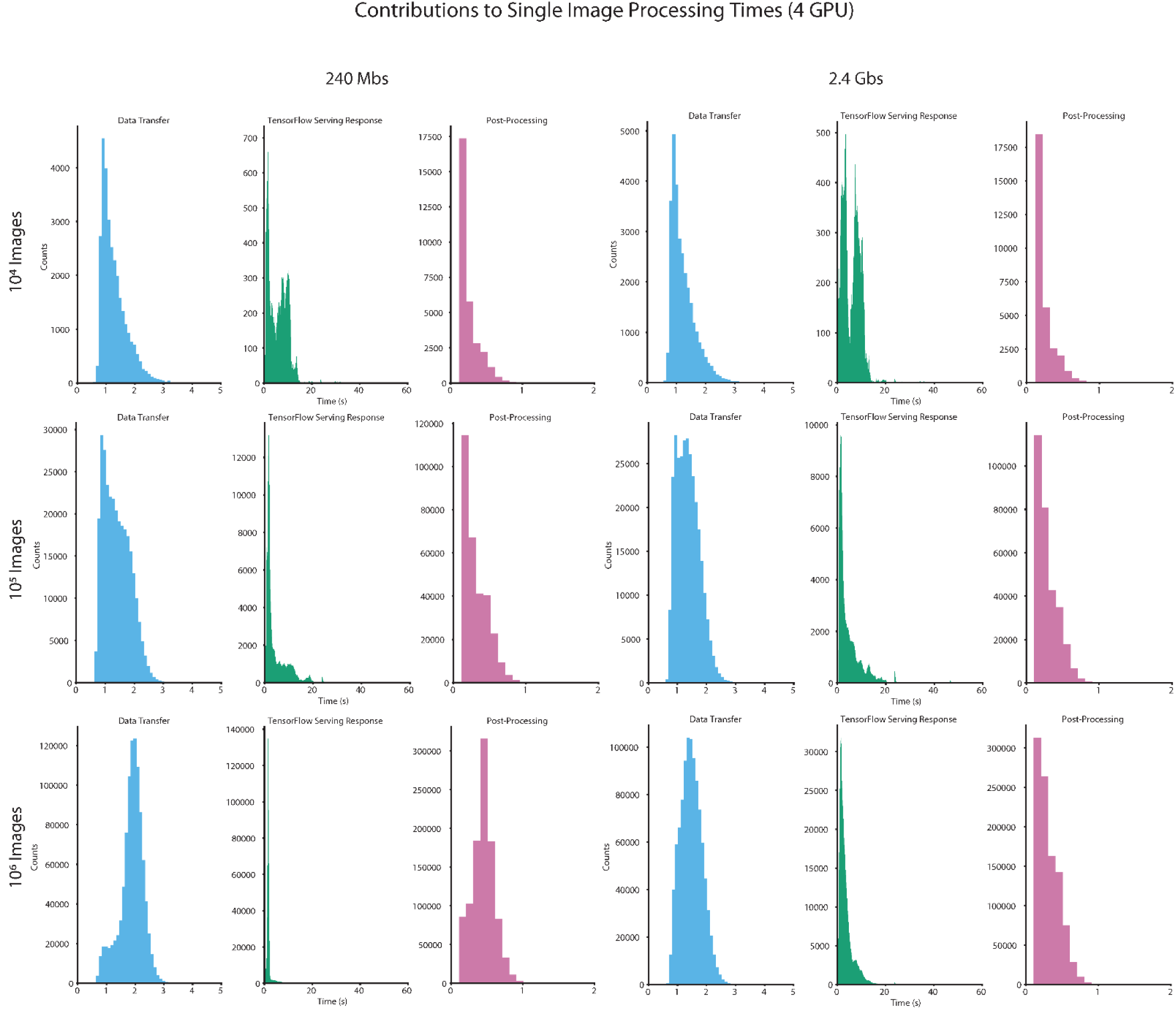
Benchmark timing data for 4 GPU clusters. Distributions of data transfer time, Tensorflow Serving response time, and postprocessing time required to process a single image.

**Figure S7:**
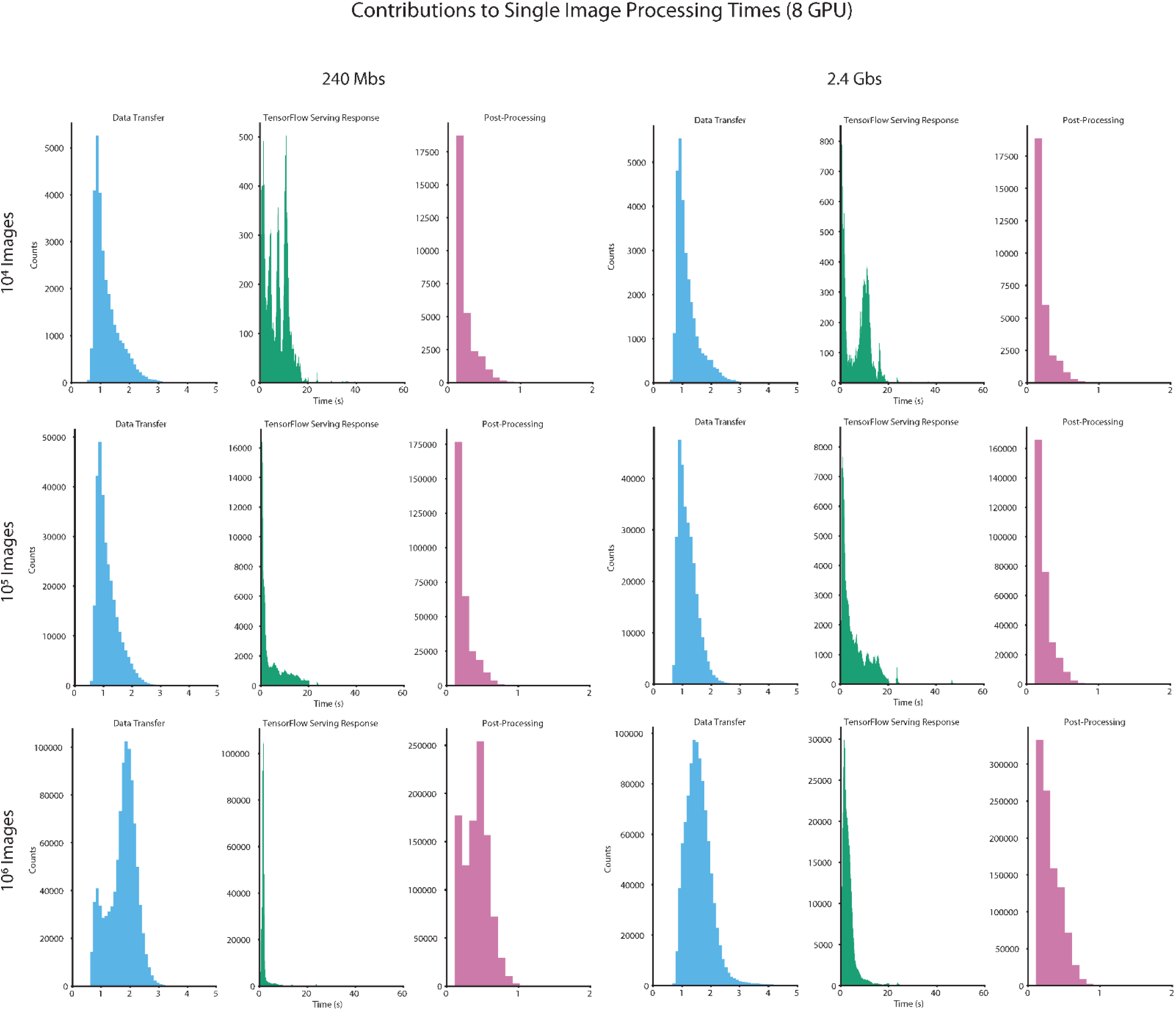
Benchmark timing data for 8 GPU clusters. Distributions of data transfer time, Tensorflow Serving response time, and postprocessing time required to process a single image.

### Data Collection and Annotation

#### Cell line acquisition and culture methods

We used the mammalian cell lines NIH-3T3, HeLa-S3, HEK 293, and RAW 264.7 to collect training data for nuclear segmentation. Genetically encoded markers (e.g., H2B-iRFP) or dyes (e.g., Hoechst) were used to label the nucleus. All cell lines were acquired from ATCC. The cells have not been authenticated and were not tested for mycoplasma contamination.

Mammalian cells were cultured in Dulbecco’s modified Eagle’s medium (DMEM, Invitrogen or Caisson) supplemented with 2 mM L-glutamine (Gibco), 100 U/mL penicillin, 100 μg/mL streptomycin (Gibco or Caisson), and 10% fetal bovine serum (FBS; Omega Scientific or Thermo Fisher) for HeLa-S3 cells or 10% calf serum (Colorado Serum Company) for NIH-3T3 cells. Cells were incubated at 37°C in a humidified 5% CO_2_ atmosphere. When 70-80% confluence was reached, cells were passaged and seeded onto fibronectin-coated glass-bottom 96-well plates (Thermo Fisher) at 10,000-20,000 cells/well. The seeded cells were incubated for 1-2h to allow cells to adhere to the bottom of the well before imaging.

#### Collection of training data

For fluorescence nuclear imaging, mammalian cells were seeded onto fibronectin-coated (Sigma, 10 μg/mL) glass-bottom 96-well plates (Nunc) and allowed to attach overnight. Growth medium was removed and replaced with imaging medium (FluoroBrite DMEM from Invitrogen; supplemented with 10 mM HEPES, 1% FBS, 2 mM L-glutamine) at least 1h prior to imaging. Cells without a genetically encoded nuclear marker were incubated with 50 ng/mL Hoechst (Sigma) prior to imaging. Cells were imaged with a Nikon Ti-E microscope or a Nikon Ti2 fluorescence microscope with environmental control (37°C, 5% CO_2_) and controlled by Micro-Manager or Nikon Elements. Images were acquired with a 20x objective (40x for RAW 264.7 cells) and with an Andor Neo 5.5 CMOS camera with 2×2 binning or a Photometrics Prime 95B CMOS camera with 2×2 binning. All data were scaled so that pixels had the same physical dimension prior to training of the deep learning model.

#### Annotation of training data

Training data were annotated on the Figure Eight platform as described previously^16^. We used both the Figure Eight annotation engine as well as our own annotation software Caliban^16^. All annotations were inspected by one of the co-authors prior to publication to ensure accuracy. Caliban is available at https://www.github.com/vanvalenlab/Caliban. Training data can be accessed through the deepcell-tf repository (https://www.github.com/vanvalenlab/deepcell-tf) under the datasets module.

### Model Architecture and Training

We use a deep watershed approach to segmentation that is inspired by previous work^17^. For each image we use a deep learning model to predict two distance maps: the distance of each cell’s pixel to the background and the distance of each cell’s pixel to the center of mass. The deep learning models used for segmentation are panoptic feature pyramids^18^ (Fig. S8). Briefly, these networks consist of a ResNet50 backbone that is connected to a feature pyramid. Prior to entering the backbone model, images are normalized (zero-meaned and then normalized by the image’s standard deviation) and concatenated with a coordinate map. We use backbone layers C3-C5 and pyramid layers P3-P7. We attach three semantic segmentation heads to the feature pyramid. The first and second heads are used to predict the inner and outer distance transforms of the instance mask image, respectively, via regression. The third head is used to predict foreground-background separation via classification.

**Figure S8:**
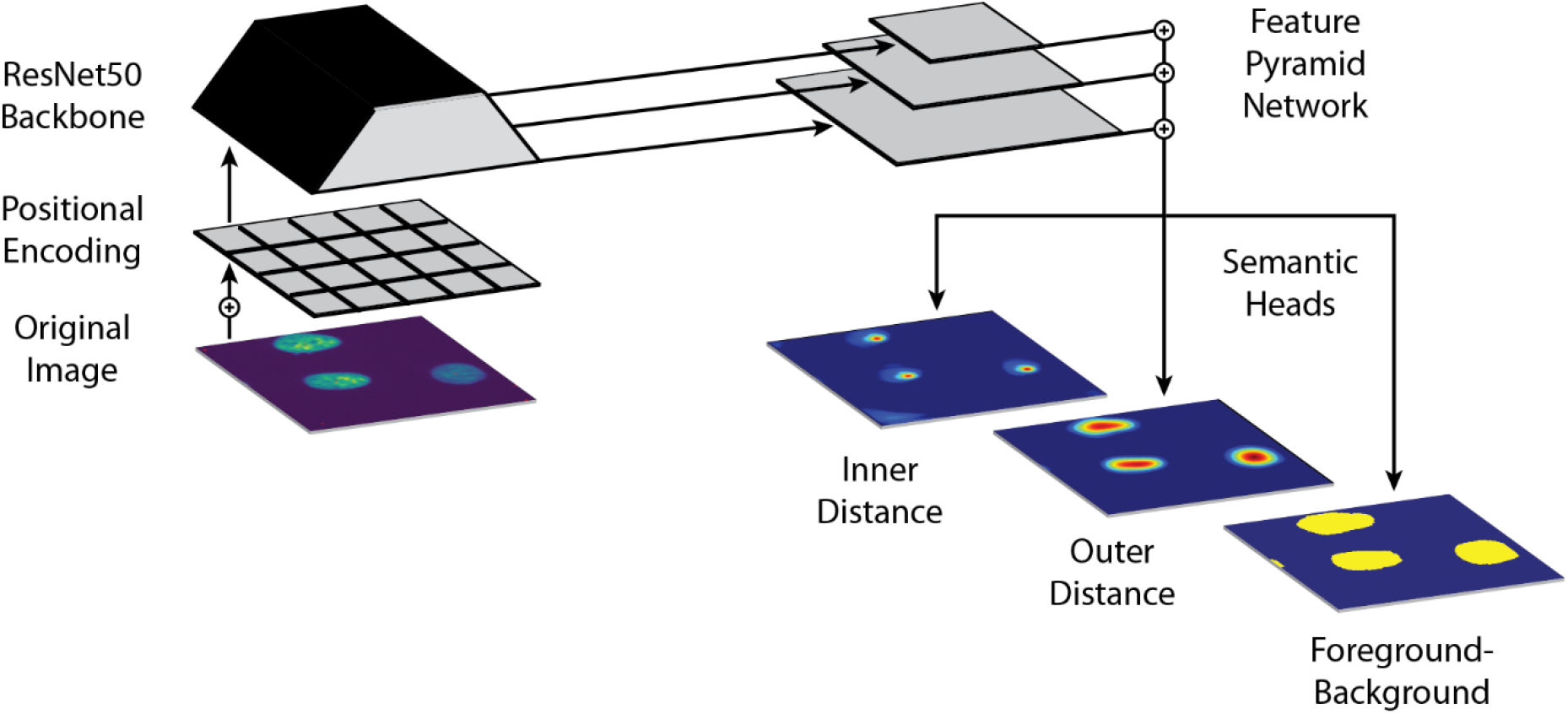
Diagram of segmentation model used for benchmarking. A ResNet50 backbone is used to extract visual features while a feature pyramid network is used to aggregate features across spatial scales. Semantic segmentation heads upsample the top of the feature pyramid to the original image size to enable prediction of distance transforms.

Models were trained with the Adam optimizer^19^ with a learning rate of 10^-5^ and clipnorm setting of 0.001. The mean squared error was used as the loss for regression tasks, while the weighted categorical cross-entropy was used for classification tasks. As described previously^17^, we weight the classification loss by 0.1 to improve the regression predictions.

Model postprocessing consisted of peak finding in the inner distance transform prediction and using the peaks to seed a marker-based watershed transform of the outer distance transform prediction. Benchmarks for models are shown in Table S1. These models were comparable to MaskR-CNN^20,21^ type models with respect to recall and precision, but are easier to deploy on large images. Code for benchmarking segmentation models is available at https://www.github.com/vanvalenlab/deepcell-toolbox.

Algorithms for cell tracking were trained as described previously^16^.

**Table S1:** Benchmarking for nuclear segmentation models. Training data were randomly divided into training (80%) and testing (20%) sets; models were trained on the training set and evaluated on the testing set. F1 scores were computed as described previously^16^.

| Model | Mean and Standard Deviation of the F1 Score for Nuclear Segmentation |
| --- | --- |
| RetinaMask<sup>21</sup> | $0.8751 \pm 0.0025$ |
| Deep Watershed | $0.8968 \pm 0.0062$ |

